# The ANTs Longitudinal Cortical Thickness Pipeline

**DOI:** 10.1101/170209

**Authors:** Nicholas J. Tustison, Andrew J. Holbrook, Brian B. Avants, Jared M. Roberts, Philip A. Cook, Zachariah M. Reagh, Jeffrey T. Duda, James R. Stone, Daniel L. Gillen, Michael A. Yassa, for the Alzheimer’s Disease Neuroimaging Initiative

## Abstract

Longitudinal studies of development and disease in the human brain have motivated the acquisition of large neuroimaging data sets and the concomitant development of robust methodological and statistical tools for quantifying neurostructural changes. Longitudinal-specific strategies for acquisition and processing have potentially significant benefits including more consistent estimates of intra-subject measurements while retaining predictive power. In this work, we introduce the open-source Advanced Normalization Tools (ANTs) registration-based cortical thickness longitudinal processing pipeline and its application to the first phase of the Alzheimer’s Disease Neuroimaging Initiative (ADNI-1) comprising over 600 subjects with multiple time points from baseline to 36 months. We demonstrate in these data that the single-subject template construction and same orientation processing results in a simultaneous minimization of residual variability and maximization of between-subject variability immediately estimable from a longitudinal mixed-effects modeling strategy. It is known from the statistical literature that optimizing these dual criteria leads to greater scientific interpretability in terms of tighter confidence intervals in calculated mean trends, smaller prediction intervals, and narrower confidence intervals for determining cross-sectional effects. This strategy is evaluated over the entire cortex, as defined by the Desikan-Killiany-Tourville labeling protocol, where comparisons are made with the cross-sectional and longitudinal FreeSurfer processing streams. Subsequent linear mixed effects modeling for identifying diagnostic groupings within the ADNI cohort is provided as supporting evidence for the utility of the proposed ANTs longitudinal framework which provides unbiased structural neuroimage processing and competitive to superior power for longitudinal structural change detection.

## 1 Introduction

Quantification of brain morphology facilitates the investigation of a wide range of neurological conditions with structural correlates, including neurodegenerative conditions such as Alzheimer’s disease [1, 2]. Essential for thickness quantification are the computational techniques which were developed to provide accurate measurements of the cerebral cortex. These include various mesh-based (e.g., [3–5]) and volumetric techniques (e.g., [6–9]).

In inferring developmental processes, many studies employ cross-sectional population sampling strategies despite the potential for confounding effects [10]. Large-scale studies involving longitudinal image acquisition of a targeted subject population, such as the Alzheimer’s Disease Neuroimaging Initiative (ADNI) [11], are designed to mitigate some of the relevant statistical issues. Analogously, much research has been devoted to exploring methodologies for properly exploiting such studies and avoiding various forms of processing bias [12]. For example, FSL’s SIENA (Structural Image Evaluation, using Normalization, of Atrophy) framework [13] for detecting atrophy between longitudinal image pairs avoids a specific type of processing bias by transforming the images to a midspace position between the two time points. As the authors point out “[i]n this way both images are subjected to a similar degree of interpolation-related blurring.” Consequences of this “interpolation-related blurring” were formally analyzed in [14] in the context of hippocampal volumetric change where it was shown that interpolation-induced artifacts can artificially create and/or inflate effect size [15]. These insights and others have since been used for making specific recommendations with respect to longitudinal image data processing [12, 16–18].

In [12, 19], the authors motivated the design and implementation of the longitudinal FreeSurfer variant inspired by these earlier insights and the overarching general principle of “treat[ing] all time points exactly the same.” It has since been augmented by integrated linear mixed effects modeling capabilities [20] and has been used in a variety of studies including pediatric cortical development [21], differential development in Alzheimer’s disease and fronto-temporal dementia [22], and fatigue in the context of multiple sclerosis [23]. Although the FreeSurfer longitudinal processing stream is perhaps one of the most well-known, other important longitudinal-specific methodologies have been proposed for characterizing cortical morphological change. Similar to FreeSurfer, cortical surfaces are generated in [24, 25] permitting vertex-wise quantitation of thickness and thickness change. Application to early infants in [24] further demonstrate the utility of targeted longitudinal considerations.

We introduced the Advanced Normalization Tools (ANTs) cortical thickness pipeline in [26] which leverages various pre-processing, registration, segmentation, and other image analysis tools that members of the ANTs and Insight Toolkit (ITK) open-source communities have developed over the years and disseminated publicly [27]. This proposed ANTs-based pipeline has since been directed at a variety of neuroimaging research topics including mild cognitive impairment and depression [28], short term memory in mild cognitive impairment [29], and aphasia [30]. Other authors have extended the general framework to non-human studies [31, 32].

In this work, we introduce the longitudinal version of the ANTs registration-based cortical thickness pipeline and demonstrate its utility on the publicly available ADNI-1 data set. In addition, we demonstrate that certain longitudinal processing choices have significant impact on measurement quality in terms of residual and between-subject variances which is known to impact the scientific interpretability of results, produce tighter confidence intervals in calculated mean trends and smaller prediction intervals, as well as less varied confidence/credible intervals for discerning cross-sectional effects. Similar to previously outlined research, we show that reorienting individual time point images to a single-subject template has favorable performance effects which guides processing choices for the proposed ANTs longitudinal pipeline. Although we limit exploration in this work to ROI-based analysis for simplifying comparison with FreeSurfer, there are several additional applications permitted by the ANTs framework such as longitudinal tensor-based morphometry, Eigenanatomy [33], and extension to non-human data.

## 2 Methods and materials

### 2.1 ADNI-1 imaging data

The strict protocol design, large-scale recruitment, and public availability of the Alzheimer’s Disease Neuroimaging Initiative (ADNI) makes it an ideal data set for evaluating the ANTs longitudinal cortical thickness pipeline. An MP-RAGE [34] sequence for 1.5 and 3.0 T was used to collect the data at the scan sites. Specific acquisition parameters for 1.5 T and 3.0 T magnets are given in Table 1 of [35]. As proposed, collection goals were 200 elderly cognitively normal subjects collected at 0, 6,12, 24, and 36 months; 400 MCI subjects at risk for AD conversion at 0, 6, 12, 18, 24, and 36 months; and 200 AD subjects at 0, 6,12, and 24 months.

**Table 1:** The 31 cortical labels (per hemisphere) of the Desikan-Killiany-Tourville atlas. The ROI abbreviations from the R brainGraph package are given in parentheses and used in later figures.

|  |  |
| --- | --- |
| 1) caudal anterior cingulate (cACC) | 17) pars orbitalis (pORB) |
| 2) caudal middle frontal (cMFG) | 18) pars triangularis (pTRI) |
| 3) cuneus (CUN) | 19) pericalcarine (periCAL) |
| 4) entorhinal (ENT) | 20) postcentral (postC) |
| 5) fusiform (FUS) | 21) posterior cingulate (PCC) |
| 6) inferior parietal (IPL) | 22) precentral (preC) |
| 7) inferior temporal (ITG) | 23) precuneus (PCUN) |
| 8) isthmus cingulate (iCC) | 24) rosterior anterior cingulate (rACC) |
| 9) lateral occipital (LOG) | 25) rostral middle frontal (rMFG) |
| 10) lateral orbitofrontal (LOF) | 26) superior frontal (SFG) |
| 11) lingual (LING) | 27) superior parietal (SPL) |
| 12) medial orbitofrontal (MOF) | 28) superior temporal (STG) |
| 13) middle temporal (MTG) | 29) supramarginal (SMAR) |
| 14) parahippocampal (PARH) | 30) transverse temporal (TT) |
| 15) paracentral (paraC) | 31) insula (INS) |
| 16) pars opercularis (pOPER) |  |

The ADNI-1 data were downloaded in May of 2014 and first processed using the ANTs cross-sectional cortical thickness pipeline [26] (4399 total images). Data was then processed using two variants of the ANTs longitudinal stream (described in the next section). In the final set of csv files (which we have made publicly available in the GitHub repository associated with this work [36]), we only included time points for which clinical scores (e.g., MMSE) were available. In total, we included 197 cognitive normals, 324 LMCI subjects, and 142 AD subjects with one or more follow-up image acquisition appointments. Further breakdown of demographic information is given in Figures 1 and 2 to provide additional perspective on the data used for this work.

**Figure 1:**
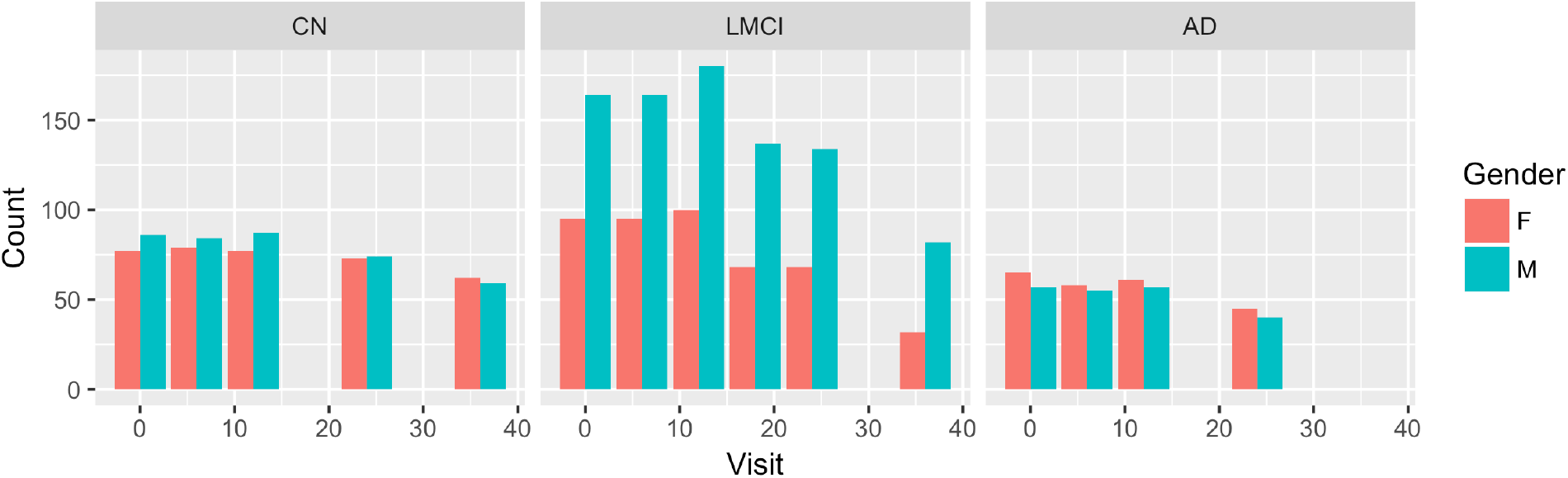
Demographic breakdown of the number of ADNI-1 subjects by diagnosis i.e., normal, mild cognitive impairment (MCI), late mild cognitive impairment (LMCI), and Alzheimer’s disease (AD). Within each panel we plot the number of subjects (by gender) per visit–baseline (“bl”) and *n* months (“m*n*”).

**Figure 2:**
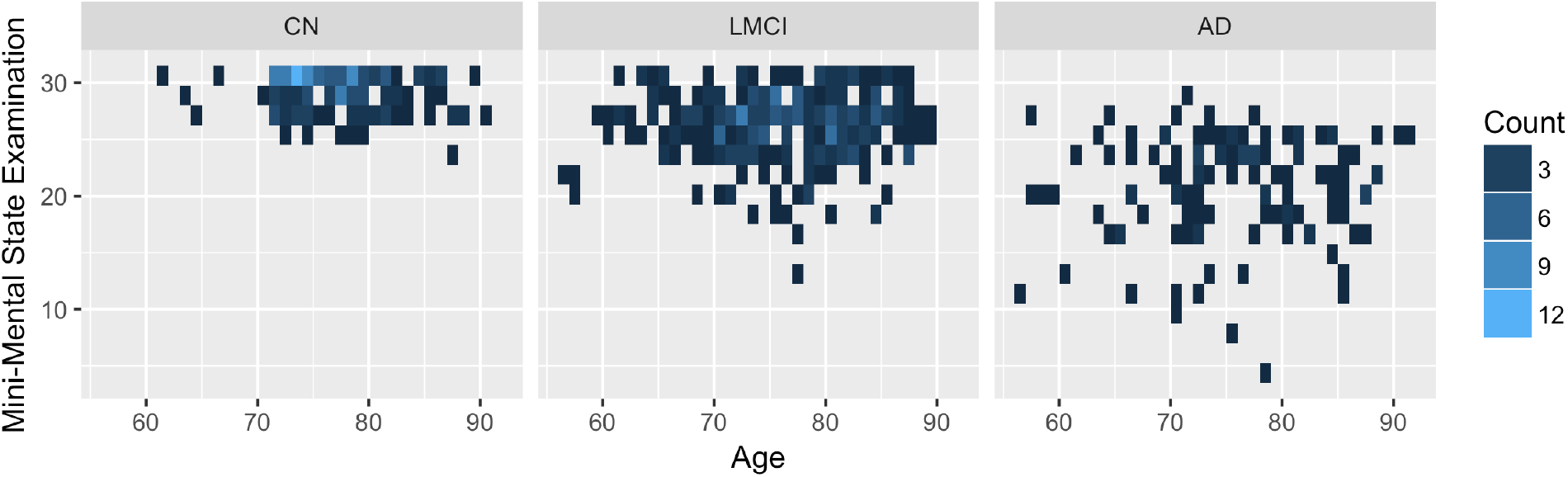
Age vs. Mini-mental examination (MMSE) scores for the ADNI-1 subjects by diagnosis providing additional demographic characterization for the subjects processed for this study.

### 2.2 ANTs cortical thickness

#### 2.2.1 Cross-sectional processing

A thorough discussion of the ANTs cross-sectional thickness estimation framework was previously provided in [26]. As a brief review, given a T1-weighted brain MR image, processing comprises the following major steps (cf Figure 1 of [26]):

1. preprocessing (e.g., N4 bias correction [37]),
2. brain extraction [38],
3. Atropos n-tissue segmentation [39], and
4. registration-based cortical thickness estimation [8].

ROI-based quantification is achieved through joint label fusion [40] of the cortex coupled with the MindBoggle-101 data. These data use the Desikan–Killiany–Tourville (DKT) labeling protocol [41] to parcellate each cortical hemisphere into 31 anatomical regions (cf Table 1). This pipeline has since been enhanced by the implementation [42] of a patch-based denoising algorithm [43] as an optional preprocessing step and multi-modal integration capabilities (e.g., joint T1- and T2-weighted image processing). All spatial normalizations are generated using the well-known Symmetric Normalization (SyN) image registration algorithm [44, 45] which forms the core of the ANTs toolkit and constitutes the principal component of ANTs-based processing and analysis.

For evaluation, voxelwise regional thickness statistics were summarized based on the DKT parcellation scheme. Test-retest error measurements were presented from a 20-cohort subset of both the OASIS [46] and MMRR [47] data sets and compared with the corresponding FreeSurfer thickness values. Further evaluation employed a training/prediction paradigm where regional cortical thickness values generated from 1205 images taken from four publicly available data sets (i.e., IXI [48], MMRR, NKI [49], and OASIS) were used to predict age and gender using linear and random forest [50] models. The resulting regional statistics (including cortical thickness, surface area [51], volumes, and Jacobian determinant values) were made available online [52]. These include the corresponding FreeSurfer measurements which are also publicly available for research inquiries (e.g., [53]). Since publication, this framework has been used in a number of studies (e.g., [54–56]).

#### 2.2.2 Unbiased longitudinal processing

Given certain practical limitations (e.g., subject recruitment and retainment), as mentioned earlier, many researchers employ cross-sectional acquisition and processing strategies for studying developmental phenomena. Longitudinal studies, on the other hand, can significantly reduce inter-subject measurement variability. The ANTs longitudinal cortical thickness pipeline extends the ANTs cortical thickness pipeline for longitudinal studies which takes into account various bias issues previously discussed in the literature [12, 14, 19].

Given *N* time-point T1-weighted MR images (and, possibly, other modalities) and representative images to create a population-specific template and related images, the longitudinal pipeline consists of the following steps:

1. (Offline): Creation of the group template and corresponding prior probability images.
2. Creation of the unbiased single-subject template (SST).
3. Application of the ANTs cross-sectional cortical thickness pipeline [26] to the SST with the group template and priors as input.
4. Creation of the SST prior probability maps.
5. (Optional): Rigid transformation of each individual time point to the SST.
6. Application of the ANTs cross-sectional cortical thickness pipeline [26], with the SST as the reference template, to each individual time-point image. Input includes the SST and the corresponding spatial priors made in Step 3.
7. Joint label fusion to determine the cortical ROIs for analysis.

An overview of these steps is provided in Figure 3 which we describe in greater detail below.

**Figure 3:**
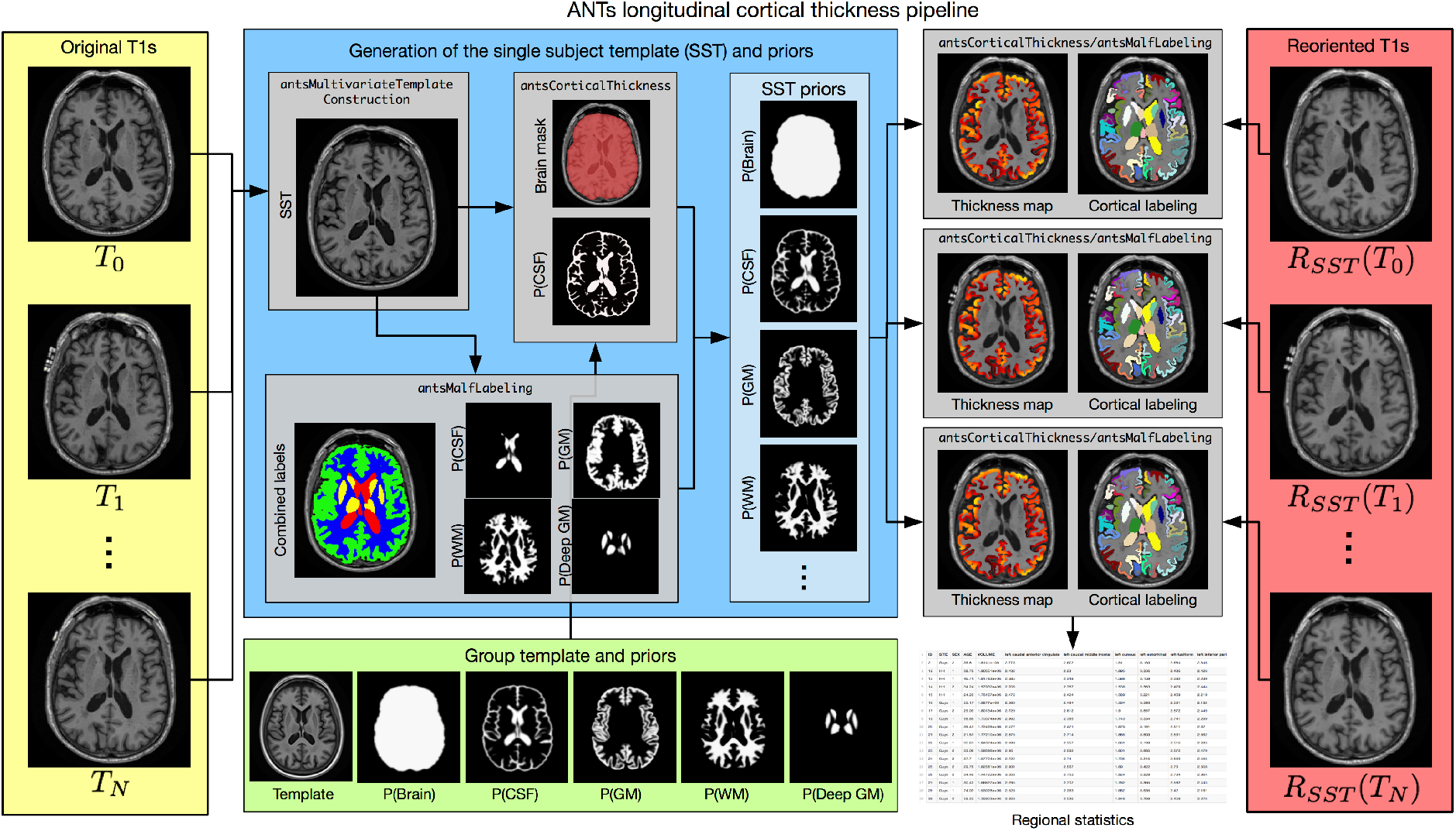
Diagrammatic illustration of the ANTs longitudinal cortical thickness pipeline for a single subject with *N* time points. From the *N* original T1-weighted images (left column, yellow panel) and the group template and priors (bottom row, green panel), the single-subject template (SST) and auxiliary prior images are created (center, blue panel). These subject-specific template and other auxiliary images are used to generate the individual time-point cortical thickness maps, in the individual time point’s native space (denoted as “ANTs Native” in the text). Optionally, one can rigidly transform the time-point images prior to segmentation and cortical thickness estimation (right column, red panel). This alternative processing scheme is referred to as “ANTs SST”. For regional thickness values, regional labels are propagated to each image using a given atlas set (with cortical labels) and joint label fusion.

##### ADNI group template, brain mask, and tissue priors

Prior to any individual subject processing, the group template is constructed from representative population data [57]. For the ADNI-1 processing described in this work, we created a population-specific template from 52 cognitively normal ADNI-1 subjects. Corresponding brain and tissue prior probability maps for the CSF, gray matter, white matter, deep gray matter, brain stem, and cerebellum were created as described in [26]. A brief overview of this process is also provided in the section concerning creation of the single-subject template. Canonical views of the ADNI-1 template and corresponding auxiliary images are given in

**Figure 4:**
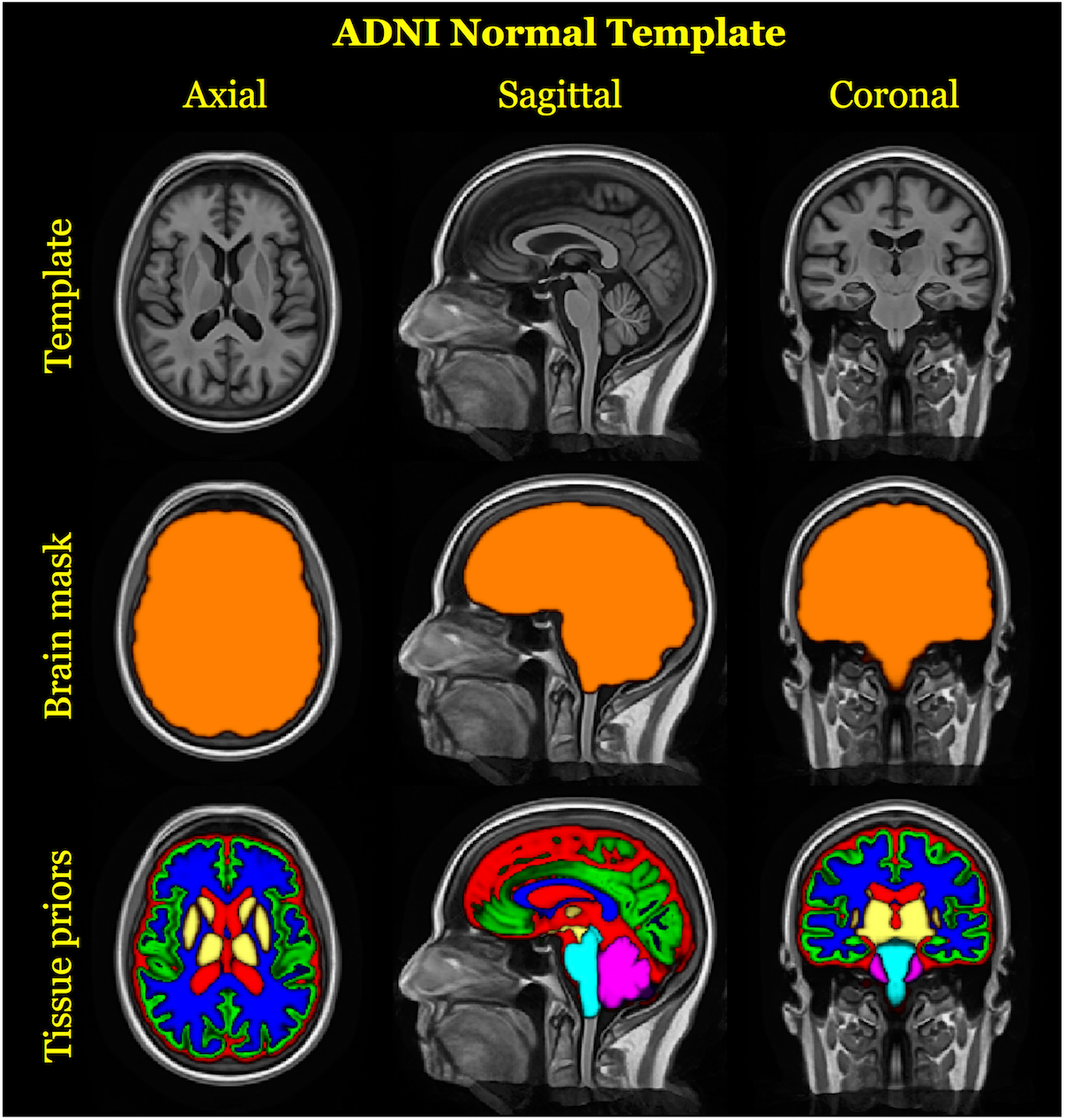
Top row: Canonical views of the template created from 52 randomly selected cognitively normal subjects of the ADNI-1 database. The prior probability mask for the whole brain (middle row) and the six tissue priors (bottom row) are used to “seed” each single-subject template for creation of a probabilistic brain mask and probabilistic tissues priors during longitudinal processing.

##### Single-subject template, brain mask, and tissue priors

With the ADNI-1 group template and prior probability images, each subject undergoes identical processing. First, an average shape and intensity single subject template (SST) is created from all time-point images using the same protocol [57] used to produce the ADNI-1 group template. Next, six probabilistic tissue maps (cerebrospinal fluid (CSF), gray matter (GM), white matter (WM), deep gray matter (striatum + thalamus), brain stem, and cerebellum) are generated in the space of the SST. This requires processing the SST through two parallel workflows. First, the SST proceeds through the standard cross-sectional ANTs cortical thickness pipeline which generates a brain extraction mask and the CSF tissue probability map, *P_Seg_*(*CSF*). Second, using a data set of 20 atlases from the OASIS data set that have been expertly annotated and made publicly available [41], a multi-atlas joint label fusion step (JLF) [40] is performed to create individualized probability maps for all six tissue types. Five of the JLF probabilistic tissue estimates (GM, WM, deep GM, brain stem, and cerebellum) and the JLF CSF estimate, *P_JLF_*(*CSF*), are used as the SST prior probabilities after smoothing with a Gaussian kernel (isotropic, *σ* = 1*mm*) whereas the CSF SST tissue probability is derived as a combination of the JLF and segmentation CSF estimates, i.e., *P*(*CSF*) = max (*P_Seg_* (*CSF*), *P_JLF_*(*CSF*)), also smoothed with the same Gaussian kernel. Finally, *P*(*CSF*) is subtracted out from the other five tissue probability maps. Note that the unique treatment of the CSF stems from the fact that the 20 expertly annotated atlases only label the ventricular CSF. Since cortical segmentation accuracy depends on consideration of the external CSF, the above protocol permits such inclusion in the SST CSF prior probability map. The final version of the SST and auxiliary images enable unbiased, non-linear mappings to the group template, subject-specific tissue segmentations, and cortical thickness maps for each time point of the original longitudinal image series.

##### Individual time point processing

The first step for subject-wise processing involves creating the SST from all the time points for that individual [57]. For the cross-sectional ANTs processing, the group template and auxiliary images are used to perform tasks such as individual brain extraction and n-tissue segmentation prior to cortical thickness estimation [26]. However, in the longitudinal variant, the SST serves this purpose. We thus deformably map the SST and its priors to the native space of each time point where individual-level segmentation and cortical thickness is estimated.

Note that this unbiased longitudinal pipeline is completely agnostic concerning ordering of the input time-point images, i.e., we “treat all time points exactly the same.”

An ANTs implementation of the denoising algorithm described in [43] is a recent addition to the toolkit and has been added as an option to both the cross-sectional and longitudinal pipelines. This denoising algorithm employs a non-local means filter [58] to account for the spatial varying noise in MR images in addition to specific consideration of the Rician noise inherent to MRI [59]. This preprocessing step has been used in a variety of imaging studies for enhancing segmentation-based protocols including hippocampal and ventricle segmentation [60], voxel-based morphometry in cannabis users [61], and anterior temporal lobe gray matter volume in bilingual adults [62]. An illustration of the transformed data resulting from image preprocessing (bias correction plus denoising) is provided in Figure 5.

**Figure 5:**
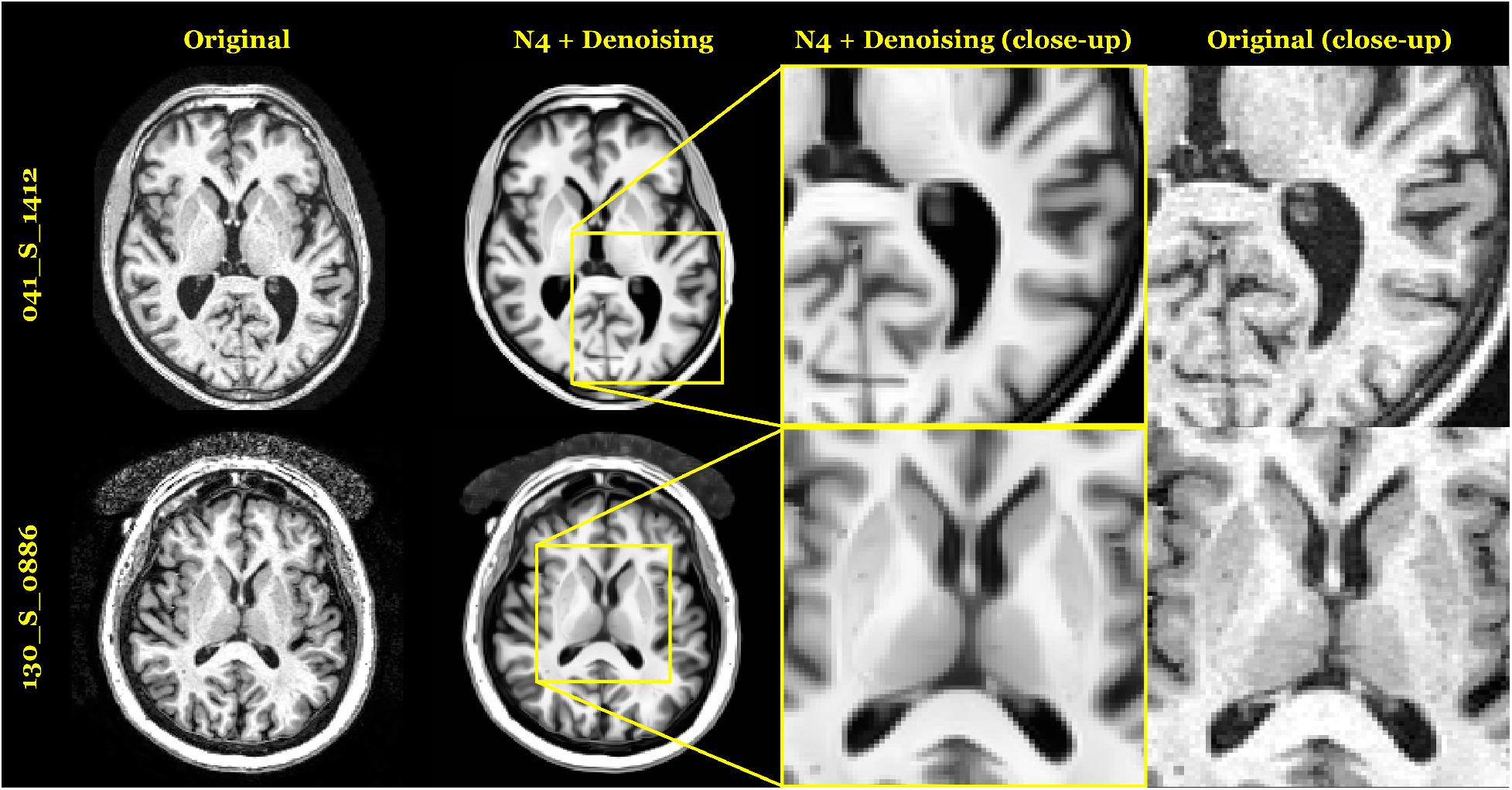
Images from two randomly chosen subjects were chosen to illustrate the effects of bias correction and denoising. The former mitigates artificial low spatial frequency modulation of intensities whereas the latter reduces the high frequency spatially-varying Rician noise characteristic of MRI.

In the FreeSurfer longitudinal stream, each time-point image is processed using the FreeSurfer crosssectional stream. The resulting processed data from all time points is then used to create a mean, or median, single-subject template. Following template creation, each time-point image is rigidly transformed to the template space where it undergoes further processing (e.g., white and pial surface deformation). This reorientation to the template space “further reduce[s] variability” and permits an “implicit vertex correspondence” across all time points [12].

The ANTs framework also permits rotation of the individual time point image data to the SST, similar to FreeSurfer, for reducing variability, minimizing or eliminating possible orientation bias, and permitting a 4-D segmentation given that the Atropos segmentation implementation is dimensionality-agnostic [39]. Regarding the 4-D brain segmentation, any possible benefit is potentially outweighed by the occurrence of “over-regularization” [12] whereby smoothing across time reduces detection ability of large time-point changes. Additionally, it is less than straightforward to accommodate irregular temporal sampling such as the acquisition schedule of the ADNI-1 protocol.

##### Registration-based cortical thickness

The underlying registration-based estimation of cortical thickness, Diffeomorphic Registration-based Estimation of Cortical Thickness (DiReCT), was introduced in [8]. Given a probabilistic estimate of the cortical gray and white matters, diffeomorphic-based image registration is used to register the white matter probability map to the combined gray/white matters probability map. The resulting mapping defines the diffeomorphic path between a point on the GM/WM interface and the GM/CSF boundary. Cortical thickness values are then assigned at each spatial location within the cortex by integrating along the diffeomorphic path starting at each GM/WM interface point and ending at the GM/CSF boundary. A more detailed explanation is provided in [8] with the actual implementation provided in the class `itk::DiReCTImageFilter` available as part of the ANTs library.

##### Joint label fusion and pseudo-geodesic for large cohort labeling

Cortical thickness ROI-based analyses are performed using joint label fusion [40] and whatever cortical parcellation scheme is deemed appropriate for the specific study. The brute force application of the joint label fusion algorithm would require *N* pairwise non-linear registrations for each time-point image where *N* is the number of atlases used. This would require a significant computational cost for a relatively large study such as ADNI. Instead, we use the “pseudo-geodesic” approach for mapping atlases to individual time point images (e.g., [63]). The transformations between the atlas and the group template are computed offline. With that set of non-linear transforms, we are able to concatenate a set of existing transforms from each atlas through the group template, to the SST, and finally to each individual time point for estimating regional labels for each image.

### 2.3 Statistical evaluation

Based on the above ANTs pipeline descriptions, there are three major variants for cortical thickness processing of longitudinal data. We denote these alternatives as:

- **ANTs Cross-sectional** (or **ANTs Cross**). Process each subject’s time point independently using the cross-sectional pipeline originally described in [26].
- **ANTs Longitudinal-SST** (or **ANTs SST**). Rigidly transform each subject to the SST and then segment and estimate cortical thickness in the space of the SST.
- **ANTs Longitudinal-native** (or **ANTs Native**). Segment and estimate cortical thickness in the native space.

For completeness, we also include a comparison with both the cross-section and longitudinal FreeSurfer v5.3 streams respectively denoted as “FreeSurfer Cross-sectional” (or “FS Cross”) and “FreeSurfer Longitudinal” (or “FS Long”).

### 2.4 Cross-sectional and longitudinal evaluation strategies

Possible evaluation strategies for cross-sectional methods have employed manual measurements in the histological [64] or virtual [65] domains but would require an inordinate labor effort for collection to be comparable with the size of data sets currently analyzed. Other quantitative measures representing “reliability,” “reproducibility,” or, more generally, “precision” can also be used to characterize such tools. For example, [66] used FreeSurfer cortical thickness measurements across image acquisition sessions to demonstrate improved reproducibility with the longitudinal stream over the cross-sectional stream. In [67] comparisons for ANTs, FreeSurfer, and the proposed method were made using the range of measurements and their correspondence to values published in the literature. However, none of these precision-type measurements, per se, indicate the utility of a pipeline-specific cortical thickness value as a potential biomarker. For example, Figure 8 in [26] confirms what was found in [67] which is that the range of ANTs cortical thickness values for a particular region exceeds those of FreeSurfer. However, for the same data, the demographic predictive capabilities of the former was superior to that of the latter. Thus, better assessment strategies are necessary for determining clinical utility. For example, the intra-class correlation (ICC) coefficient used in [26] demonstrated similarity in both ANTs and FreeSurfer for repeated acquisitions despite the variance discrepancy between both sets of measurements. This is understood with the realization that the ICC takes into account both inter-observer and intra-observer variability.

Similarly, evaluation strategies for longitudinal studies have been proposed with resemblance to those employed for cross-sectional data such as the use of visual assessment [24], scan-rescan data [12, 25], and 2-D comparisons of post mortem images and corresponding MRI [25]. In addition, longitudinal methods offer potential for other types of assessments such as the use of simulated data (e.g., atrophy [12, 25], infant development [24]) where “ground-truth” is known and regression analysis of longitudinal trajectories of cortical thickness [68].

### 2.5 Cortical residual and between-subject thickness variability

For a longitudinal biomarker to be effective at classifying subpopulations, it should have low residual variation and high between-subject variation. Without these simultaneous conditions, subpopulation distinctions would not be possible (e.g., if measurements within the subject vary more than those between subjects). A summary measure related to the ICC statistic [69] is used to quantify this intuition for assessing relative performance of these cross-sectional and longitudinal ANTs pipeline variants along with the cross-sectional and longitudinal FreeSurfer streams. Specifically, we use longitudinal mixed-effects (LME) modeling to quantify pipeline-specific between-subject and residual variabilities where comparative performance is determined by maximizing the ratio between the former and the latter. Such a quantity implies greater within-subject reproducibility while distinguishing between patient sub-populations. As such this amounts to higher precision when cortical thickness is used as a predictor variable or model covariate in statistical analyses upstream.

LME models comprise a well-established and widely used class of regression models designed to estimate cross-sectional and longitudinal linear associations between quantities while accounting for subject-specific trends. As such, these models are useful for the analysis of longitudinally collected cohort data. Indeed, [20] provides an introduction to the mixed-effects methodology in the context of longitudinal neuroimaging data and compare it empirically to competing methods such as repeated measures ANOVA. For more complete treatments of the subject matter, see [69] and [70]. LME models are also useful for estimating and comparing residual and between-subject variability after conditioning out systematic time trends in longitudinally measured data. In the context of the current investigation, by fitting LME models to the data resulting from cross-sectional and longitudinal processing techniques, we are able to quantify the relative performance of each approach with respect to residual, between-subject, and total variability in a way that [71] hint at in their exposition of the longitudinal FreeSurfer stream.

As previously noted we observed a longitudinal sampling of cortical thickness measurements from the 62 parcellated cortical DKT regions. To assess the above variability-based criteria while accounting for changes that may occur through the passage of time, we used a Bayesian LME model for parameter estimation. Let 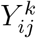 denote the *i^th^* individual’s cortical thickness measurement corresponding to the *k^th^* region of interest at the time point indexed by *j*. Under the Bayesian paradigm we utilized a model of the form

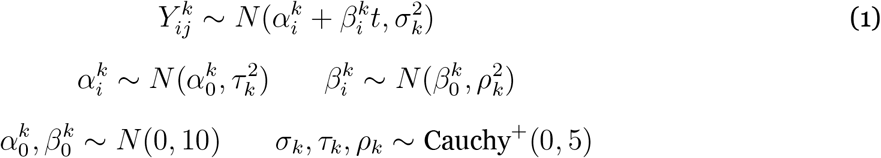

where specification of variance priors to half-Cauchy distributions reflects commonly accepted best practice in the context of hierarchical models [72]. These priors concentrate mass near zero but have heavy tails, meaning small variance values are expected but large variance values are not prohibited. Even so, results demonstrated robustness to parameter selection.

In Model (1), *τ_k_* represents the between-subject standard deviation, and *σ_k_* represents the within-subject standard deviation, conditional upon time, and 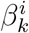 denotes the subject-specific slopes of cortical thickness change. For each region *k*, the quantity of interest is thus the ratio

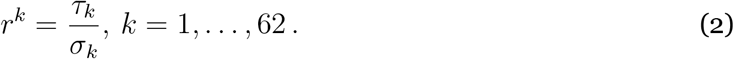

The posterior distribution of *r^k^* was summarized via the posterior median where the posterior distributions were obtained using the Stan probabilistic programming language [73]. The R interface to Stan was used to calculate the point estimates of Model (1) for cortical thickness across the different pipelines using the default parameters. The csv files containing the regional cortical thickness values for all five pipelines, the Stan model file, and the R script to run the analysis and produce the plots are all located in the GitHub repository created for this work [36].

This ratio is at the heart of classical statistical discrimination methods as it features both in the ANOVA methodology and in Fisher’s linear discriminant analysis. These connections are important since the utility of cortical thickness as a biomarker lies in the ability to discriminate between patient sub-populations with respect to clinical outcomes. In particular, [74] (Sections 9.6.2 and 9.6.5) demonstrate the role that randomness and measurement error in explanatory variables play in statistical inference. When the explanatory variable is fixed but measured with error (as is plausible for cortical thickness measurements), the residual variance divided by the between subject variance is proportional to the bias of the estimated linear coefficient when the outcome of interest is regressed over the explanatory variable (Example 9.2). In short, the larger the *r^k^*, the less bias in statistical analyses. When the explanatory variable is considered random and is measured with error (a common assumption in the measurement error literature [75, 76]), this bias is expressed as attenuation of regression coefficient estimates to zero by a multiplicative factor *r^k^*/(1 + *r^k^*) (Example 9.3). Thus, larger *r^k^* means less less attenuation bias and hence more discriminative capacity. Note that effect estimator bias is not the only problem–the residual variance is increased by a factor proportional to *r^k^*/(1 + *r^k^*) ([74], Chapter 3). The same authors refer to the combination of bias and added variance as a ‘double whammy’. Indeed, a worse reliability ratio causes greater bias in multiple linear regression in the presence of collinearity and even biases the estimators for other covariates, progression through time included (cf [76], Section 3.3.1). The same authors state that this bias is typical even in generalized linear models (Section 3.6) and use the ratio as a measure of reliability even in the longitudinal context (Section 11.9).

### 2.6 Regional diagnostic contrasts based on cortical atrophy

The variance ratio explored in the previous section is a generic desideratum for statistical assessment of performance over the set of possible use cases. In this section, we narrow the focus to the unique demographical characteristics of the ADNI-1 study data and look at performance of the various pipelines in distinguishing between diagnostic groups on a region-by-region basis. Previous work has explored various aspects of Alzheimer’s disease with respect to its spatial distribution and the regional onset of cerebral atrophy. For example, although much work points to the entorhinal cortex as the site for initial deposition of amyloid and tau [77], other evidence points to the basal forebrain as preceding cortical spread [78]. Other considerations include the use of hippocampal atrophy rates as an image-based biomarker of cognitive decline [79], differentiation from other dementia manifestations (e.g., posterior cortical atrophy [80]), and the use of FreeSurfer for monitoring disease progression [81]. Thus, longitudinal measurements have immediate application in Alzheimer’s disease research. To showcase the utility of the ANTs framework, we compare the generated longitudinal measurements and their ability to differentiate diagnostic groups (i.e., CN vs. LMCI vs. AD).

Pipeline-specific LME models were constructed for each DKT region relating the change in cortical thickness to diagnosis and other pertinent covariates. In the notation of [82], these regional LME models are defined as:

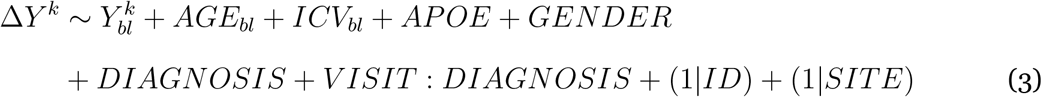

where Δ*Y^k^* is the change in thickness of the *k^th^* DKT region from baseline (bl) thickness measurement 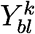. We also include random intercepts for both the individual subject (ID) and the acquisition site. Modeling was performed in R using the `lme4` package [83] followed by Tukey post-hoc analyses with false discovery rate (FDR) adjustment using the `multcomp` package in Rto test the significance of the “LMCI–CN”, “AD–LMCI”, and “AD–CN” diagnostic contrasts.

## 3 Results

All imaging data were processed through the five processing streams (i.e., FS Cross, FS Long, ANTs Cross, ANTs SST, and ANTs Native) on the high performance computing cluster at the University of California, Irvine (UCI). Based on the evaluation design described in the previous section, we compare pipeline performance when applied to the ADNI-1 data. Specifically, we calculate the variance ratios, described in Section 2.5, for each of the 62 DKT regions for each of the five pipelines. These are compared and discussed. We then explore how this general criterion for evaluating measurement quality applies specifically to a longitudinal analysis of the ADNI-1 data in discriminating the diagnostic stages of Alzheimer’s disease.

After processing the image data through the various pipelines, we tabulated the regional thickness values and made them available as .csv files online in the corresonding GitHub repository [36]. We also provide the Perl scripts used to run the pipelines on the UCI cluster and the R scripts used to do the analysis below. Additional figures and plots have also been created which were not included in this work. For example, spaghetti plots showing absolute thickness and longitudinal thickness changes are contained in the subdirectory CrossLong/Data/RegionalThicknessSpaghettiPlots/.

### 3.1 Cortical residual and between-subject thickness variability

The LME model defined in (1) was used to quantify the between-subject and residual variance with the expectation that maximizing the former while minimizing the latter optimizes measurement quality in terms of prediction and confidence intervals. Figure 6 provides the resulting 95% credible intervals for the distributions of region-specific variance ratios *r^k^* = *τ_k_*/*σ_k_* for each of the five pipelines. Based on the discussion in the previous section, superior methododologies are designated by larger variance ratios.

**Figure 6:**
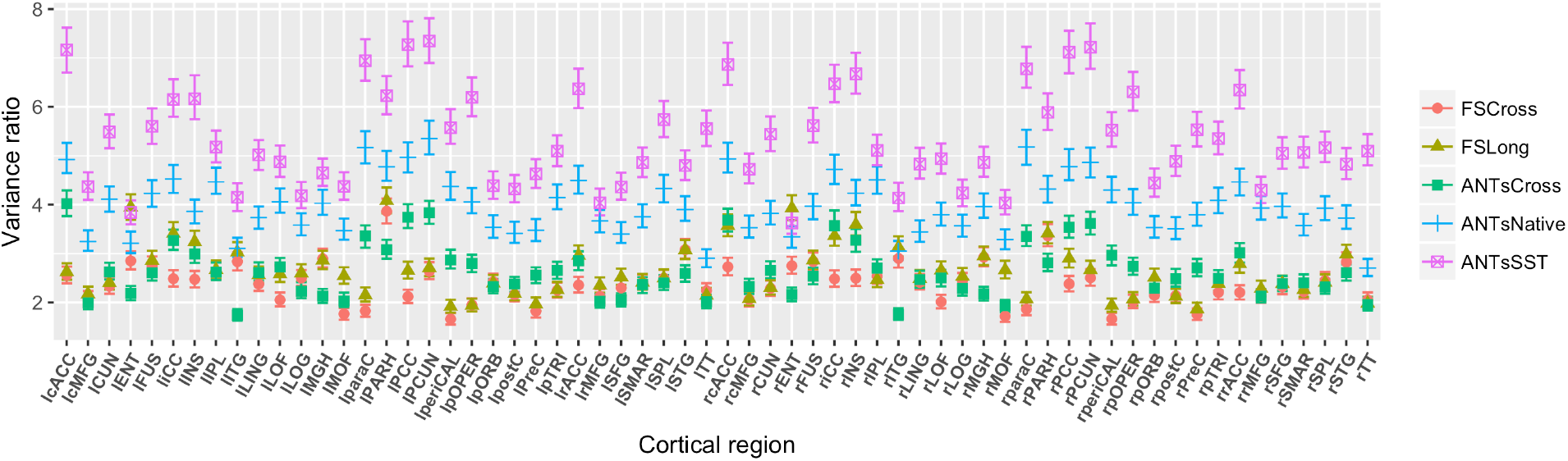
95% credible intervals of the region-specific variance ratios *r^k^* = *τ_k_*/*σ_k_* are presented for each processing method. The ANTs SST method dominates the others across the majority of regions—its point estimates (posterior medians) are greater than those of the other processing methods except for the left and right EC values in FreeSurfer Long (although there is significant overlap in the credible intervals in those regions). These results also suggest that longitudinal processing is to be preferred for both packages.

ANTs SST has the highest ratio variance across most of the 62 regions over the other methods. It rarely overlaps with ANTs Native and never with ANTs Cross. In contrast to the majority of FreeSurfer regional ratio variances (from both FS Cross and FS Long) which are smaller than those of the ANTs pipelines, FS Long has larger ratio values for the EC region with the only overlap in the credible intervals with ANTs SST.

The plot in Figure 7 shows a relative summary of all the regional quantities for all three variance measurements (residual, between-subject, and variance ratio) via box plots. These relative distributions show that both between-subject and residual quantities contribute to the disparities in the ratio evaluation metric. Finally, we overlay the variance ratio values on the corresponding regions of a 3-D rendering of the ADNI template (Figure 8) to provide an additional visual comparison between the methods.

**Figure 7:**
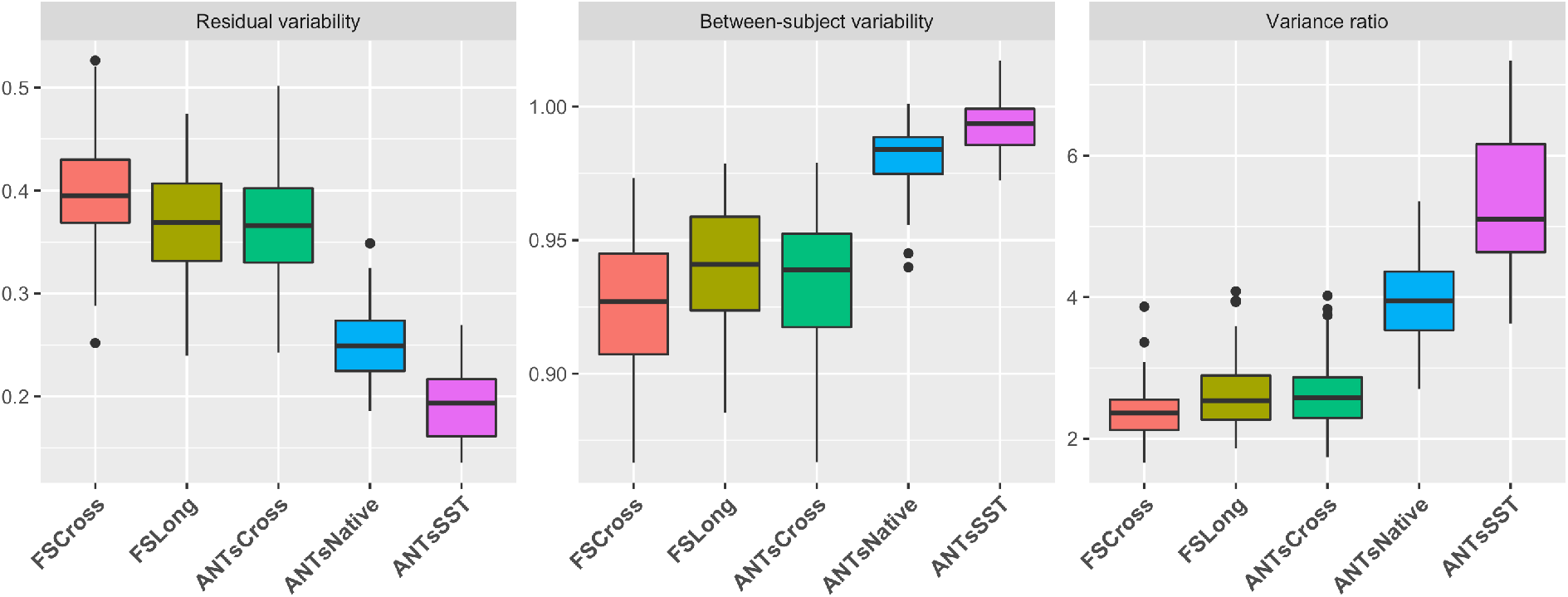
Box plots showing the distribution of the residual variability, between subject variability, and ratio of the between-subject variability and residual variability for each of the 62 DKT regions. Note that the “better” measurement maximizes this latter ratio.

**Figure 8:**
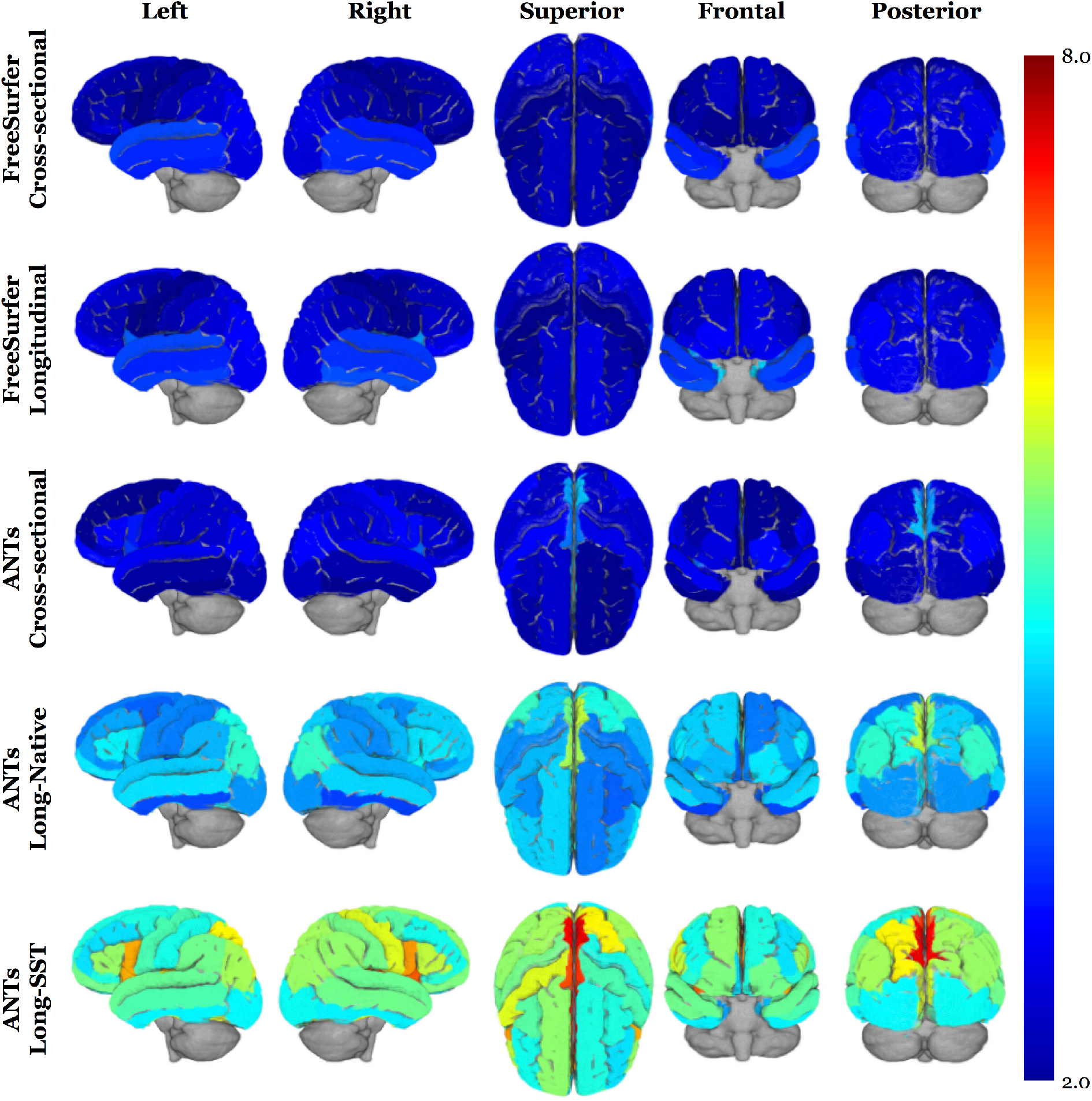
3-D volumetric rendering of the regional variance ratio values on the generated ADNI template. The higher variance ratios indicate greater between-subject to residual variability.

### 3.2 Regional diagnostic contrasts based on cortical atrophy

The LME model described in Equation (3) was used to determine region-by-region contrasts for each pairing “LMCI–CN”, “AD–LMCI”, and “AD–CN” using post-hoc Tukey significance testing. It should be noted that no subjects were included that switched diagnostic groups during the acquired study schedule. These findings are provided in Tables 2 and 3. The adjusted *p*-values were log-scaled for use in specifying the individual color cell for facilitating visual differentiation. Each cell contains the corresponding 95% confidence intervals. Figure 9 provides a side-by-side comparison of the distribution of log-scaled *p*-values separated into left and right hemispherical components and grouped according to contrast.

**Table 2:**
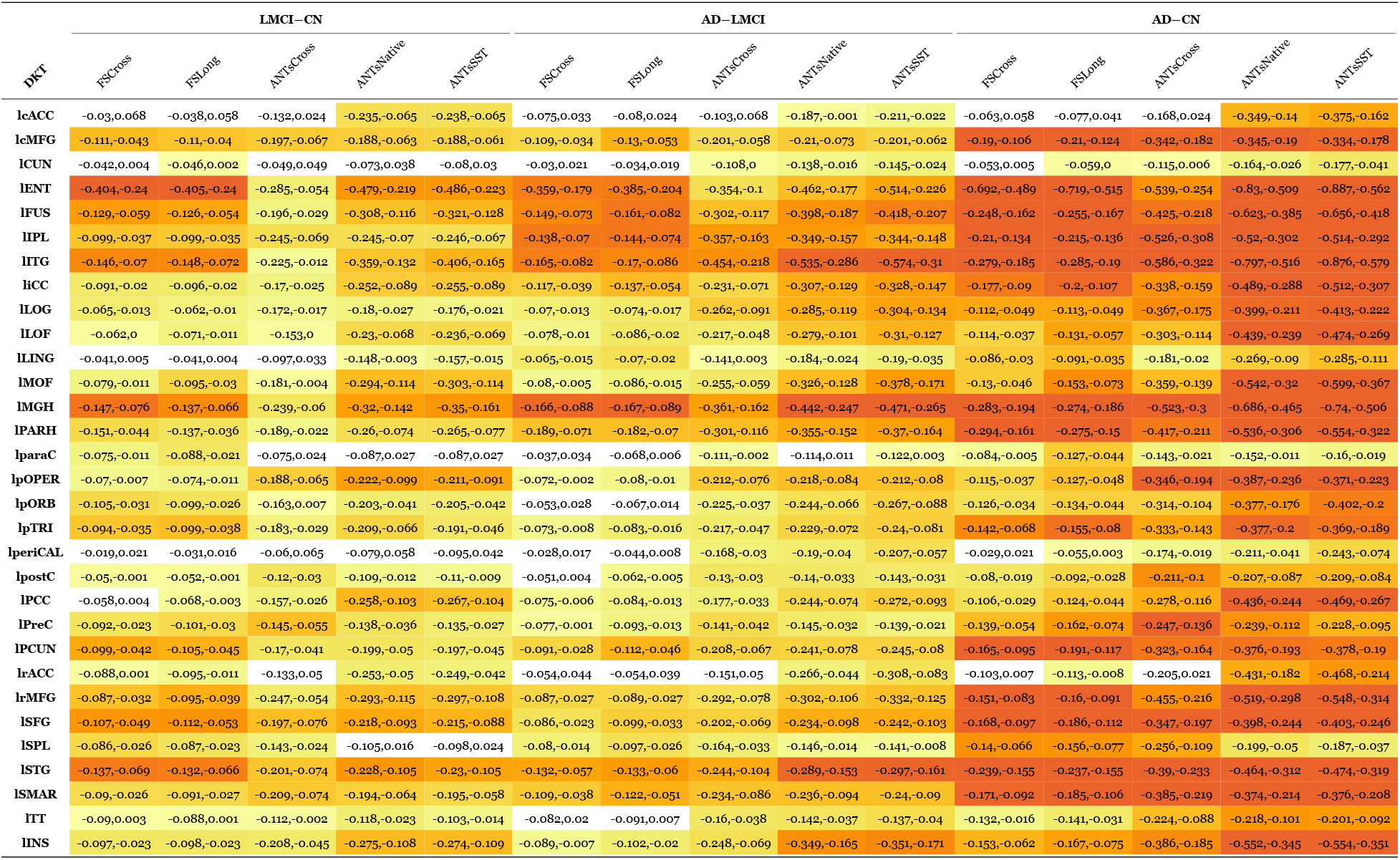
95% confidence intervals for the difference in slope values for the three diagnoses (CN, LMCI, AD) of the ADNI-1 data set for each DKT region of the left hemisphere. Each cell is color-coded based on the adjusted log-scaled *p*-value significance from dark orange (*p* < 1e-10) to yellow (*p* = 0.1). Absence of color denotes nonsignificance.

**Table 3:**
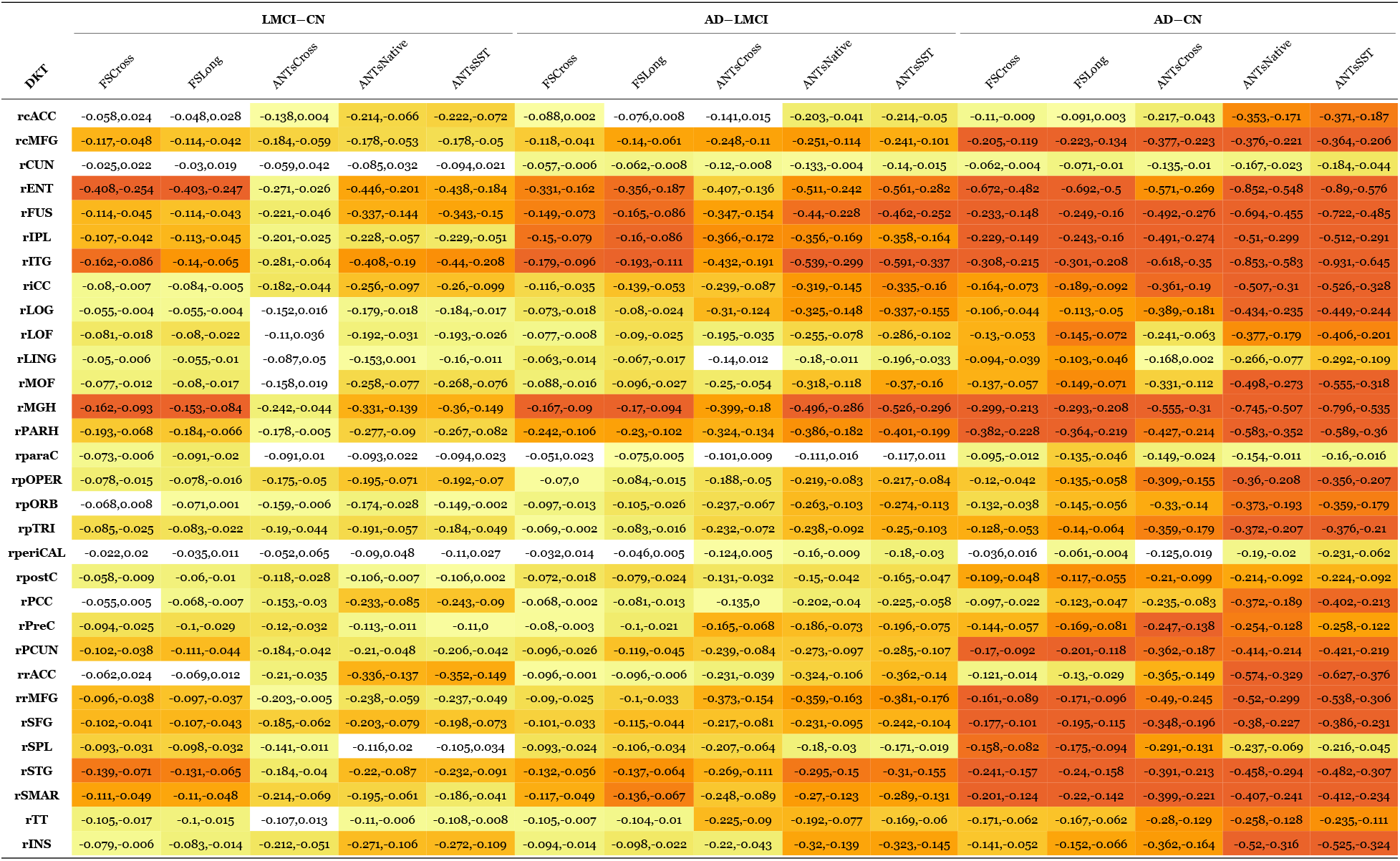
95% confidence intervals for the difference in slope values for the three diagnoses (CN, LMCI, AD) of the ADNI-1 data set for each DKT region of the right hemisphere. Each cell is color-coded based on the adjusted log-scaled *p*-value significance from dark orange (*p* < 1e-10) to yellow (*p* = 0.1). Absence of color denotes nonsignificance.

**Figure 9:**
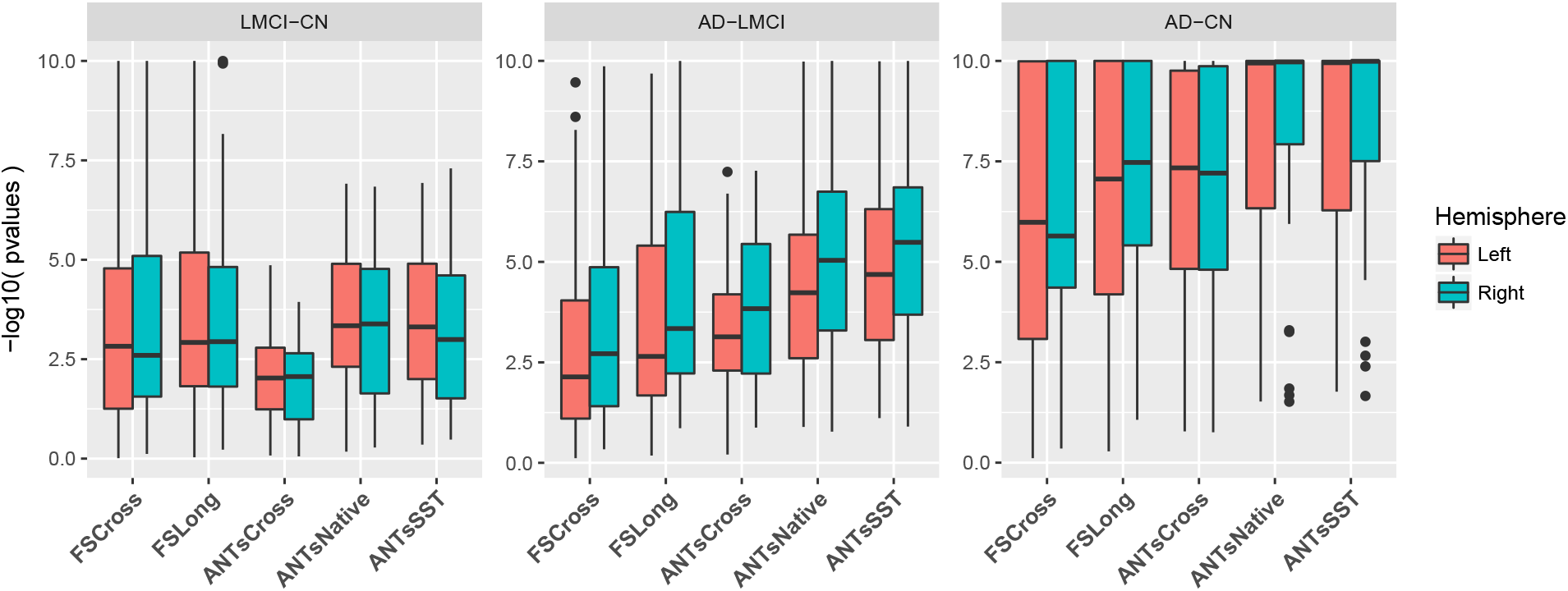
Log-scaled p-values summarizing Tables 2 and 3 demonstrating performance differences across cross-sectional and longitudinal pipelines for the three diagnostic contrasts.

Consideration of performance over all three diagnostic pairings illustrates the superiority of the longitudinal ANTs methodologies over their ANTs cross-sectional counterpart. Several regions demonstrating statistically significant non-zero atrophy in ANTs Native and ANTs SST do not manifest similar trends in ANTs Cross (e.g., LMCI–CN: lateral occipital gyri). Pronounced differences between the ANTs longitudinal vs. cross-sectional methodologies can be seen in both the LMCI–CN and AD–CN contrasts. Although ANTs Cross demonstrates discriminative capabilities throughout the cortex and, specifically, in certain AD salient regions, such as the entorhinal and parahippocampal cortices, the contrast is not nearly as strong as the other methods including FS Cross and FS Long thus motivating the use of longitudinal considerations for processing of AD data.

Differentiation between the longitudinal methods is not as obvious although trends certainly exist. In general, for differentiating CN vs. LMCI, all methods are comparable except for ANTs Cross. However, for the other two diagnostic contrasts “AD–LMCI” and “AD–CN”, the trend is similar to what we found in the evaluation via the variance ratio, viz., the longitudinal ANTs methods tend towards greater contrast means versus ANTs Cross and the two FreeSurfer methods. Looking at specific cortical areas, though, we see that comparable regions (“comparable” in terms of variance ratio) are consistent with previous findings. For example, we noted in the last section that FSLong has a relatively large variance ratio in the entorhinal regions which is consistent with the results seen in Tables 2 and 3.

## 4 Discussion

Herein we detailed the ANTs registration-based longitudinal cortical thickness framework which is designed to take advantage of longitudinal data acquisition protocols. This framework has been publicly available as open-source in the ANTs GitHub repository for some time. It has been employed in various neuroimaging studies and this work constitutes a formalized exploration of performance for future reference. It inherits the performance capabilities of the ANTs cross-sectional pipeline providing high reliability for large studies, robust registration and segmentation in human lifespan data, and accurate processing in data (human and animal) which exhibit large shape variation. In addition, the ANTs longitudinal pipeline accounts for the various bias issues that have been associated with processing such data. For example, denoising and N4 bias correction mitigate the effects of noise and intensity artifacts across scanners and visits. The use of the single-subject template provides an unbiased subject-specific reference space and a consistent intermediate space for normalization between the group template and individual time points. Undergirding all normalization components is the well-performing SyN registration algorithm which has demonstrated superior performance for a variety of neuroimaging applications (e.g., [63, 84]) and provides accurate correspondence estimation even in the presence of large anatomical variation. Also, given that the entire pipeline is image-based, conversion issues between surface- and voxel-based representations [85] are non-existent which enhances inclusion of other imaging data and employment of other image-specific tools for multi-modal studies (e.g., tensor-based morphometry, longitudinal cortical labeling using joint label fusion and the composition of transformations). All ANTs components are built from the Insight Toolkit which leverages the open-source developer community from academic and industrial institutions leading to a robust (e.g., low failure rate) software platform which can run on a variety of platforms.

With respect to these data and AD in general, the ANTs longitudinal cortical thickness pipelines uses unbiased diffeomorphic registration to provide robust mapping of individual brains to group template space and, simultaneously, high-resolution sensitivity to subtle longitudinal changes over time. Both advantages are relevant to AD. High baseline atrophy levels in AD lead to the need for robustness to large deformations. Sensitivity to subtle longitudinal change over time is particularly relevant to early or preclinical AD studies due to the relatively reduced atrophy rates and smaller difference from control populations. We demonstrate that our approach leads to competitive or superior estimates of annualized atrophy that are biologically plausible in AD populations and that may, in the future, support the use of T1 neuroimaging to detect treatment effects in clinical trials. Furthermore, in ADNI-1, we report a zero percent failure rate with no subject-specific tuning required.

Over 600 subjects from the well-known longitudinal ADNI-1 data set with diagnoses distributed between cognitively normal, LMCI, and AD were processed through the original ANTs cross-sectional framework [26] and two longitudinal variants. One of the variants, ANTs SST, is similar to the FreeSurfer longitudinal stream in that each time-point image is reoriented to an unbiased single-subject template for subsequent processing. ANTs Native, in contrast, estimates cortical thickness in the native space while also using tissue prior probabilities generated from the SST.

Comparative assessment utilized LME models to determine the between-subject to residual variance ratios over the 62 regions of the brain defined by the DKT parcellation scheme where higher values indicate greater generic statistical salience. In these terms, ANTs SST outperformed all other pipeline variants including both the FreeSurfer longitudinal and cross-sectional streams. Regional disparities between the ANTs SST and Native pipelines point to increases in both between-subject and residual variances which might be due to reorientation to a common space similar to other longitudinal strategies.

Further evidence motivating the longitudinal strategies proposed in this work and elsewhere stems from the subsequent exploration of differentiating between diagnostic groups using LMEs with the change in cortical thickness as an outcome variable. Almost across the entire cortex, longitudinal strategies (both ANTs and FreeSurfer) outperformed their cross-sectional counterparts in pairwise differentiation of diagnostic groups although these trends varied based on region and diagnosis. In the context of AD, where certain regions have increased saliency in terms of neuroscientific research, and practical considerations might give more weight to certain diagnostic results over others, further exploration is required to tease out these subtle differences and their implications for future research.

One interesting finding was the performance of FS Long in the EC regions where the variance ratios was slightly larger than those of ANTs Long/Native where the credible intervals have significant overlap. Given the small volume and indistinguishability from surrounding structures, segmentation of the EC can be relatively difficult [86]. This segmentation complexity has led to EC-specific [87] and related [88] strategies for targeted regional processing. For this work, we wanted to avoid such tuning and simply employ off-the-shelf input parameters and data. Future work will explore refining input template priors in these problematic regions for ANTs-based estimation of cortical thickness.

These findings promote longitudinal analysis considerations and motivates such techniques over cross-sectional methods for longitudinal data despite the increase in computational costs. While we focus on cortical thickness in this work, there are obvious limitations with the ANTs volume-based framework. Without a direct reconstruction of the cortical surfaces, many important cortical properties (e.g., surface area, cortical folding, sulcal depth, and gyrification) [89] cannot be generated in a straightforward manner. Additional work will want to examine these features more closely working towards a more comprehensive idea of how structure changes. This will help determine the relative importance of such cortical features and will undoubtedly guide future methodological development.

However despite these deficiencies, being inherently voxel-based, the ANTs framework does have advantages not explored in this work but certainly to be utilized in future research. Specifically, the voxel-based input/output processing is conducive to voxel-based analysis strategies (e.g., Eigenanatomy [33]) and straightforward application to non-human research domains. Also, tensor-based morphometric data are directly extracted from the output of the longitudinal processing. And while mesh-based geometric measures are unavailable, digital analogs (e.g., surface area from the digitized Crofton formula [51] and surface curvature [90]) provide a convenient data format for integrated data analysis. Finally, given the importance of structural data, such as T1-weighted images, for other types of neuroimaging studies (e.g., resting state fMRI and diffusion tensor imaging), the longitudinal processing stream provides convenient output for facilitating these other types of analyses.

The ANTs longitudinal pipeline provides several additional features that may be worth investigation in future studies. The segmentation approach provides tissue probability maps that may be used in identifying abnormalities of white matter or in voxel-wise studies of gray matter density. The longitudinal formulation of the pipeline is also likely to improve the variance ratio for other transformation-based measurements such as the log-jacobian, often employed in tensor-based morphometry [91]. Local folding and other curvature-based metrics are available, as well, through ANTsR [92]. These quantification tools, individually or jointly, may provide insight into aging and neurodegeneration and will be the subject of future evaluation efforts.

The longitudinal thickness framework is available in script form within the ANTs software library along with the requisite processing components (cf Appendix). All generated data used for input, such as the ADNI template and tissue priors, are available upon request. As previously mentioned, we also make available the csv files containing the regional thickness values for all three pipelines.

## 5 Appendix

### 5.1 Implementation overview

The script `antsLongitudinalCorticalThickness.sh` performs cortical thickness estimation for a longitudinal image series from a single subject. The following principal steps are performed:

1. A single-subject template (SST) is created from all the time point images.
2. The tissue prior probability images are generated for the SST. These six tissues are label 1: CSF, label 2: cortical gray matter, label 3: white matter, label 4: deep gray matter, label 5: brain stem, and label 6: cerebellum. Prior probability creation involves the following steps:

1. The SST is passed through `antsCorticalThicknes.sh`.
2. The brain extraction posterior for the SST is created by smoothing the brain extraction mask created during 2a.
3. If labeled atlases are not provided, we smooth the posteriors from 2.1 to create the SST segmentation priors, otherwise we use the `antsJointFusion` program to create a set of posteriors using the script `antsCookTemplatePriors.sh`.
3. Using the SST + priors, each subject is processed through the `antsCorticalThickness.shscript`.

A typical command line call is:

~~~
antsLongitudinalCorticalThickness.sh \
-d ${imageDimension} \
-e ${brainTemplate} \
-m ${brainExtractionProbabilityMask} \
-p ${brainSegmentationPriors}
-o ${outputPrefix}
${anatomicalImages[@]}
~~~

### 5.2 Input parameters

- `imageDimension`: dimensionality of the input images. Can handle 2 or 3 dimensions.
- `brainTemplate`: the group template. We have made several publicly available along with the prior tissue and brain extraction images (https://figshare.com/articles/ANTs_ANTsR_Brain_Templates/915436).
- `brainExtractionProbabilityMask`: prior probability image for the whole brain corresponding to the `brainTemplate`.
- `brainSegmentationPriors`: prior probability images for the six brain tissues mentioned above. These files are specified with the relevant labels, e.g., `prior1.nii.gz`, `prior2.nii.gz,prior3.nii.gz, prior4.nii.gz, prior5.nii.gz`, and `prior6.nii.gz`. The command line argument is specified in C-style formatting, e.g., `prior%d.nii.gz`.
- `anatomicalImages`: the time point images for a single subjects.
- other optional input parameters are available. `antsLongitudinalCorticalThickness -h` provides a listing of the full set of parameters, their descriptions, and other help information.

### 5.3 Output

In the specified output directory, the following subdirectories are created:

~~~
• ${outputPrefix}SST
• ${outputPrefix}${anatomicalImagesPrefix[0]}
• ${outputPrefix}${anatomicalImagesPrefix[1]}
• ${outputPrefix}${anatomicalImagesPrefix[2]}
• …
~~~

Each subdirectory contains the output of `antsCorticalThickness.sh` applied to the corresponding image. Output consists of the following files:

- `BrainExtractionMask`: Brain extraction mask in subject space.
- `BrainNormalizedToTemplate`: Extracted brain image normalized to the template space.
- `BrainSegmentation0N4`: Input to the segmentation algorithm. It is not brain ex-tracted, but is bias-corrected. If multiple images are used for segmentation, there will be `BrainSegmentation1N4` and so on. The brain extracted version of this is `ExtractedBrain0N4`.
- `BrainSegmentation`: Segmentation image, one label per tissue class. The number of classes is determined by the input priors.
- `BrainSegmentationPosteriors1`: Posterior probability of class 1. A similar image is produced for all classes. The numbering scheme matches the input priors.
- `CorticalThickness`: Cortical thickness image in subject space.
- `CorticalThicknessNormalizedToTemplate`: Cortical thickness image in template space.
- `ExtractedBrain0N4`: Brain-extracted version of BrainSegmentation0N4.
- `SubjectToTemplate1Warp, SubjectToTemplate0GenericAffine.mat`: Transforms to be used when warping images from the subject space to the template space.
- `SubjectToTemplateLogJacobian`: Log of the determinant of the Jacobian, quantifies volume changes in the subject to template warp.
- `TemplateToSubject0Warp, TemplateToSubject1GenericAffine.mat`: Transforms to be used when warping images from the template to the subject space.

In addition to these files, the SST subdirectory contains additional warps, suffixed `SubjectToGroupTemplateWarp and SubjectToTemplate0GenericAffine.mat`, that can be used to warp each time point image to the group template. These are a combination of the subject to SST warp, and the SST to group template warp. Also included are the SST brain and tissue prior probability images.

## Acknowledgments

Additional support to N.T. and M.Y. provided by NIMH R01 MH102392 and NIA R21 AG049220, P50 AG16573.

Data collection and sharing for this project was funded by the Alzheimer’s Disease Neuroimaging Initiative (ADNI) (National Institutes of Health Grant U01 AG024904) and DOD ADNI (Department of Defense award number W81XWH-12-2-0012). ADNI is funded by the National Institute on Aging, the National Institute of Biomedical Imaging and Bioengineering, and through generous contributions from the following: AbbVie, Alzheimer’s Association; Alzheimer’s Drug Discovery Foundation; Araclon Biotech; BioClinica, Inc.; Biogen; Bristol-Myers Squibb Company; CereSpir, Inc.; Cogstate; Eisai Inc.; Elan Pharmaceuticals, Inc.; Eli Lilly and Company; EuroImmun; F. Hoffmann-La Roche Ltd and its affiliated company Genentech, Inc.; Fujirebio; GE Healthcare; IXICO Ltd.; Janssen Alzheimer Immunotherapy Research & Development, LLC.; Johnson & Johnson Pharmaceutical Research & Development LLC.; Lumosity; Lundbeck; Merck & Co., Inc.; Meso Scale Diagnostics, LLC.; NeuroRx Research; Neurotrack Technologies; Novartis Pharmaceuticals Corporation; Pfizer Inc.; Piramal Imaging; Servier; Takeda Pharmaceutical Company; and Transition Therapeutics. The Canadian Institutes of Health Research is providing funds to support ADNI clinical sites in Canada. Private sector contributions are facilitated by the Foundation for the National Institutes of Health (www.fnih.org). The grantee organization is the Northern California Institute for Research and Education, and the study is coordinated by the Alzheimer’s Therapeutic Research Institute at the University of Southern California. ADNI data are disseminated by the Laboratory for Neuro Imaging at the University of Southern California.

